# Endoplasmic Reticulum Alterations in Living Cells Expressing INF2 Variants Causing Glomerulosclerosis and Peripheral Neuropathy

**DOI:** 10.1101/2024.05.18.594805

**Authors:** Quynh Thuy Huong Tran, Naoyuki Kondo, Hiroko Ueda, Yoshiyuki Matsuo, Hiroyasu Tsukaguchi

**Author notes:** Corresponding authors: Hiroyasu Tsukaguchi, MD, PhD Second Department of Internal Medicine, Division of Nephrology, Kansai Medical University 2-5-1 Shinmachi Hirakata, Osaka, 573-1191, Japan.

## Abstract

The endoplasmic reticulum (ER) is a single dynamic, and continuous network, which mediates a variety of biological processes. The ER spreads throughout the cytoplasm as an interconnected network mainly consisting of flat (sheets) or reticular (tubules) networks. Such ER integrity is regulated by actin-microtubule interaction. INF2 is an actin assembly factor that exclusively located in ER and is mutated in hereditary form of glomerulopathy (focal segmental glomerulosclerosis, FSGS) and peripheral neuropathy (Charcot-Marie Tooth, CMT-DIE, MIM 614455). It remains unclear how INF2 variants could affect ER morphology.

High-resolution, live-imaging of HeLa cells revealed that pathogenic INF2-CAAX variants disrupt peripheral ER complexity, generating focal clustering of polygonal tubules and preferential sheet-like appearance. G73D (causing CMT+FSGS) induced more remarkable alterations than T161N, N202S and R218W (leading to FSGS). Both actin and microtubule inhibitors shifted the ER balance towards sheet predominance and focal compaction of ER tubules, suggesting the role of cytoskeleton in shaping tubular ER networks. INF2 variants induced mitochondria fragmentation with peripheral mis-distribution. The mitochondrial alterations correlated with the degree of cytoskeletal disorganization, leading to defective respiratory function. Moreover, lysosomal trafficking was restricted by INF2 variants in the cell cortex. These organelle-cytoskeletal interactions were more remarkably impaired by CMT+FSGS variant than in FSGS variants.

Our observations underscore that INF2 variants disrupt ER integrity by disorganizing cytoskeletons, which leads to defective mitochondria function and vesicle trafficking in INF2 disorders. INF2 CMT+FSGS variants impair ER-organelles interaction more prominent than FSGS variants, suggesting the existence of specific mediators for CMT+FSGS variants.

## INTRODUCTION

The endoplasmic reticulum (ER) comprises a continuous network of peripheral tubules, perinuclear sheets or matrix^1–4^. Tubule and sheet conformation transitions dynamically each other, depending upon interaction with the ER shaping proteins as well as cytoskeletons^5^. Peripheral ER is composed of polygon structure linked by three-way junctions (TWJ) in a highly dynamic manner. Cluster of TWJs forms the ER matrices^1^ (Fig 1B, 1C). Proper ER morphology is important to maintain function of highly specialized cells like podocyte and neurons. ER intergrity is critical for various cellular processes, Ca homeostasis, lipid synthesis, and organelle trafficking^4^. Dynamic actin filaments serve multiple cellular functions, such migration, morphogenesis, endocytosis, and organelle trafficking^6,7^. The organization of filamentous actin (F-actin) networks is governed by actin regulatory proteins, including formins, which facilitate F-actin assembly and remodeling^8,9^. Among the formin members, Diaphanous-related formins, such as mDia and INF2, have been extensively studied. Both are regulated by autoinhibition through self-binding between DID and DAD, diaphanous inhibitor and autoregulatory domains, respectively (Sup Fig S1A)^10,11^. There are two INF2 isoforms of CAAX or non-CAAX and the former predominates in the kidney (Fig 1A). INF2 has a unique feature that mediates both F-actin polymerization and depolymerization (Sup Fig S1B)^12,13^. The ER form INF2-CAAX is involved in mitochondria fission at the ER-mitochondria contacts^6,10,14,15^. INF2 influences microtubule (MT) stability and distribution^16,17^. The roles of ER dysregulation have been extensively studied in human neuro-degenerated disorders^3^ but the relevance in other cell lineages remain investigated.

**Figure 1.**
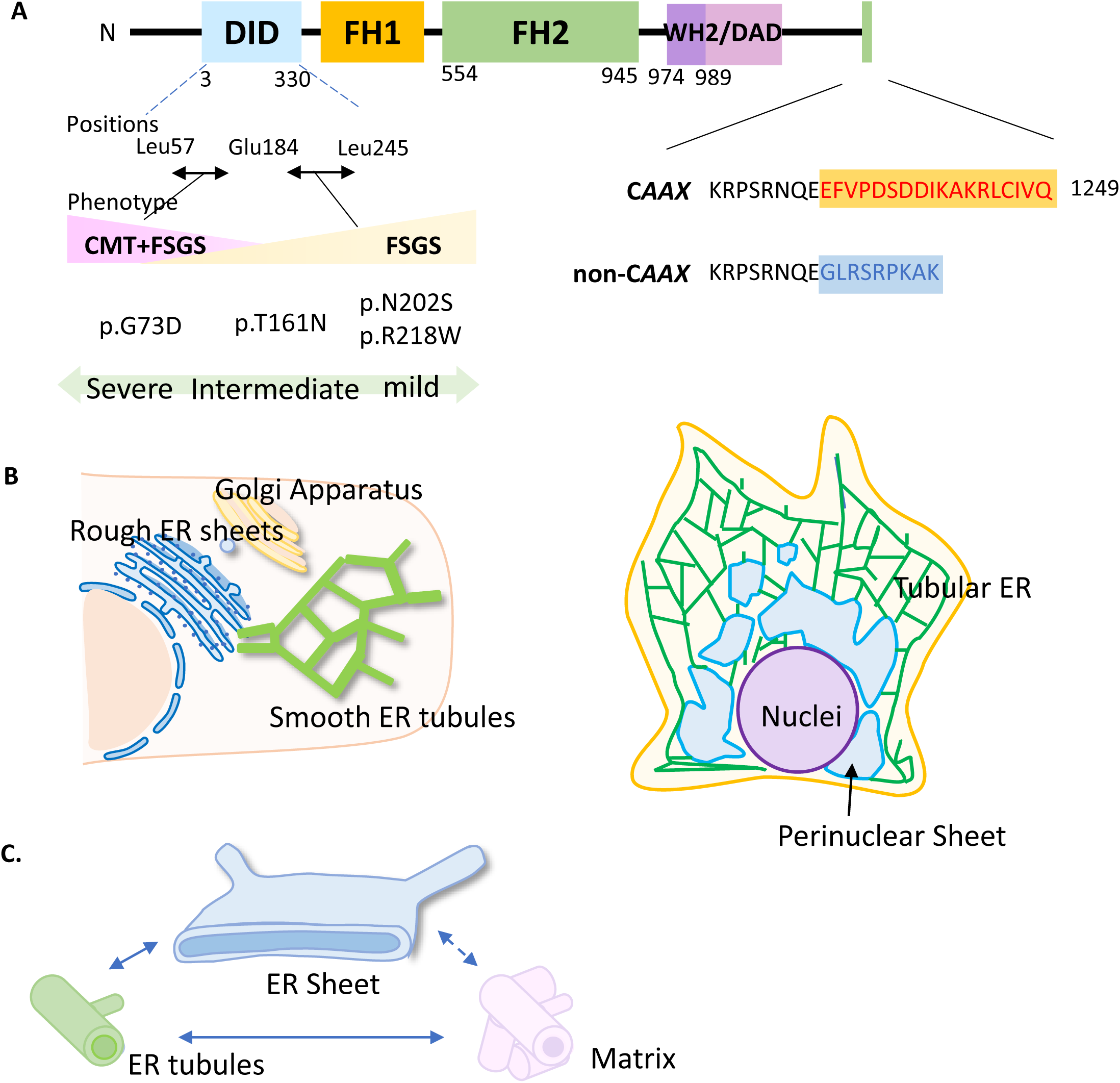
Domain structure and locations of disease variants in human *INF2*. **A. Domain structure of human INF2 and locations of variants.** Human *INF2* is multidomain and homo-dimeric protein that mainly consists of two major components: one is a regulatory domain that comprise a DID and DAD, the other is an actin-organizing unit of FH1 and FH2 domains. The amino acid numbering is shown below each box. There are two isoforms of human INF2-*CAAX* (ER-resident) and INF2-non-C*AAX* (cytoplasmic form)^10,26^. The INF2-*CAAX* predominate in the kidney^30^. *INF2* variants exclusively cluster in the DID domain: CMT+FSGS variants locate in the N-terminal half of DID Leu57-Glu183, whereas FSGS variants reside in the C-terminal half of DID Glu184-Leu245. DID: Diaphanous Inhibitory Domain. DAD: Diaphanous Autoregulatory Domain. FH1: Formin Homology 1. FH2: Formin Homology 2. **B. Schematic diagram showing the domain organization of the ER, with tubular and sheet morphologies.** ER membrane is enriched in the perinuclear region (rough ER), whereas pripheral ER is composed of tubular polygon (smooth ER). **C. Subdomains of the peripeharl ER.** The three components constitute the peripheral ER are shown. A tubule-to-sheet conformation (arrow) is dynamically regulated by the ER proteins as well as actin-microtubule networks. Cluster of tubules forms the Matrix^1-6^. The transition between sheet and matrices remain elusive (dot arrow).

Missense variants in INF2-DID domain were first identified as a gene for familial focal segmental glomerulosclerosis (FSGS)^18,19^. INF2 mutations have also been implicated in Charcot-Marie-Tooth disease with concurrent occurrence of FSGS (CMTDIE, MIM 614455), a dual phenotype affecting both motor and sensory neuropathies^20^. Expression studies with INF2 in mammalian cells showed that INF2 is implicated in the cytoskeletal integrity that mediate the cell shape, polarity and organelle dynamics^14,21–23^. However, detailed mechanisms by which INF2 regulates ER morphology and organelle contacts in concert with cytoskeleton remain explored.

The present study is aimed to evaluate the effects of INF2 variants on ER-cytoskeletons and ER-organelles interaction by use of a high-resolution microscopy and to compare the properties between CMT+FSGS and FSGS variants in living transfected HeLa cells. Our previous study with fixed cells revealed that CMT+FSGS variants cause severe cellular damage, whereas FSGS variants give relatively milder insults^23^. The fixed cell studies may have missed dynamic cytoskeleton-organelle interactions because the fine and transient architectures might attenuate during fixation. The spinning-disk confocal microscopy visualizes a small (∼70 nm) and fast-moving (5ms) vesicle with minimum photobleaching damage. Here we investigated the impact of INF2 variants on the cytoskeleton-organelle interactions of living cells, particularly focusing on ER morphology and ER-organelle interface. We found that the pathogenic INF2 variants disintegrate a tubule-to-sheet balance and expansion of peripheral ER by altering coordinated interaction between actin and microtubules. Our observations revealed that the defective ER morphology may be related to a variety of organelle defects affecting mitochondria positions and functions, and lysosomal vesicle trafficking.

## METHOD

### Plasmids, markers for cytoskeleton and organelles

Expression constructs of INF2 variants R218W, N202S, T161N and G73D, which were identified in our cohort, were generated by the gene synthesis (Genscript)^23^ based upon the original human wild-type (WT) INF2 construct (NM_022489, isoform 1, CAAX form, N-terminal eGFP tagged in pcDNA3.1 vector). The organelle markers used in the study are shown in the Supplementary Methods.

### Actin and microtubule inhibitors

To test the responses to the actin inhibitor, cells were incubated with Cytochalasin D (Fujifilm, 034-25881, 1µM) for 30 minutes and were observed at 2h after switching to the CytoD-free fresh medium. For the microtubule inhibitor studies, we used the identical incubation time and observation procedure with Nocodazole (M1404, Sigma-Aldrich, 2.5 µg/ml).

Real-time imaging of filopodia in live cells was obtained by capturing snapshots for at least 20 cells per variant using a Dragon-Fly spinning-disk confocal microscopy. Time-lapse images (minimum 10-time points) were imported into Fiji, stacked, and converted to 8-bit images. Areas with irrelevant signals was removed. The Filopodyan plugin in Fiji automatically analyze shape and movement changes. Segmentation parameters were set constant for both WT and INF2 variants: Threshold = Huang, ED Iterations = 20, LoG sigma = 3. Analysis was performed on at least 20 cells per variant.

### Cell Culture and Transfection

The human cervical cancer (HeLa) (RCB0007) or monkey kidney (COS-7) (RCB0539) were obtained from Riken and cultured in D-MEM supplemented with 10% Fetal Bovine Serum (26140-079, Gibco) at 37°C. Sub-confluent HeLa cells, which underwent less than 20 passages, were used. The transfection methods and imaging manipulation in this study are shown in Supplementary Methods

### Mitochondrial Respiration Assay

HeLa cells were transfected with eGFP-tagged WT-INF2, T161N, and G73D variants. After 12 hours, mitochondrial function was analyzed by measuring the Oxygen Consumption Rate using a Seahorse XFp Analyzer (Agilent). Three reagents were used: Oligomycin (1µM) to block ATP synthase, FCCP (1µM) to dissipate the proton gradient, and antimycin A and rotenone (0.5 µM each) to inhibit complexes III and I, respectively. Transfection efficiency was normalized by staining cells with Hoechst 33342 and visualizing GFP signals using Bz-X810 microscope (Keyence). Efficiency was calculated as the frequency of GFP-positive cells among total DAPI-positive cells. Normalized data were analyzed using WAVE software to compare differences between wild-type INF2 and variants (T161N, G73D).

## RESULTS

### Genetic features of pathogenic INF2 variants in single FSGS and dual CMT+FSGS disease

We previously identified the total 10 pathogenic, heterozygous, missense INF2 variants in our FSGS cohort: six displayed single FSGS phenotype while four exhibited dual CMT+FSGS manifestations^23^. Among these, we studied two mild (causing FSGS alone: R218W, N202S), one intermediate (FSGS alone: T161N) and one severe (CMT+FSGS: G73D) variants (Fig 1, Supplementary Method).

### Colocalization of GFP-*INF2* and ER marker

To investigate the effects of INF2 variants on the peripheral ER morphology, we initially assessed INF2 colocalization with the luminal ER marker, calreticulin^24^. In WT-INF2 expressing cells, INF2 mostly colocalized within the ER compartments labelled by Calreticulin, generating a reticular pattern with perinuclear ER cisterna. INF2 punctuates often existed in vicinity of TWJs of tubular polygons (Fig 2). Both R218W and T161N variants caused focal clustering of the peripheral tubular ER network, kinetically favoring more sheet or matrix pattern. In contrast, G73D variants occasionally retained within swollen ER tubules, displaying a coarse granular pattern (ER fragmentation) in addition to the sheet or matrix accumulation (Fig 3, Sup Fig S2). Our observations indicate that INF2 variants mislocate and accumulate in disintegrated ER compartments compared to the WT-INF2 (Sup Fig S3).

**Figure 2.**
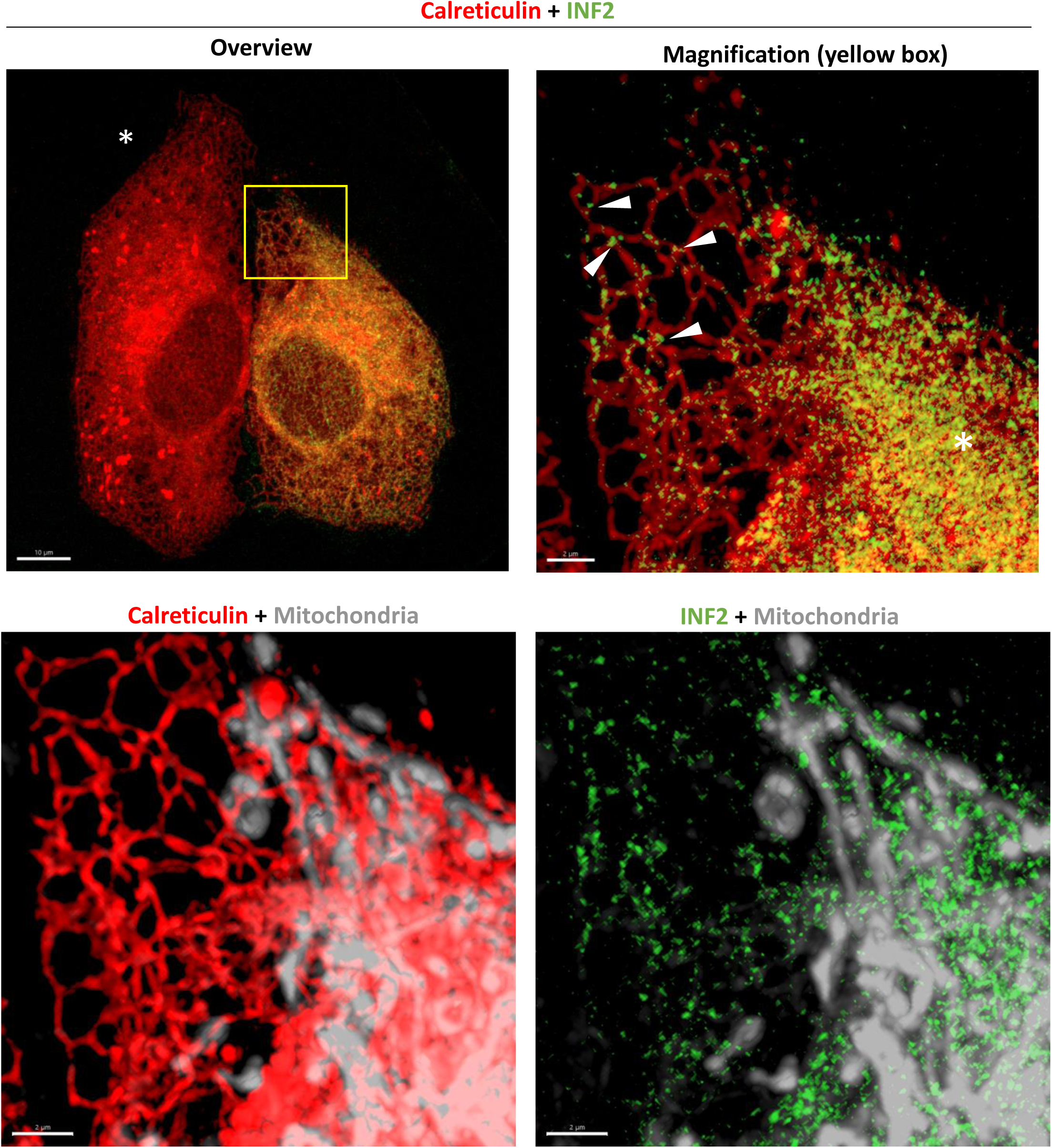
High-resolution images of ER and mitochondria morphology in living HeLa cells expressing wild-type INF2. GFP tagged, wild-type INF2 (WT, CAAX isoform) (green) are transiently co-transfected with an ER-marker mCherry-Calreticulin (red) in HeLa cells. After co-labelling with mitotracker, high-resolution images are captured by a DragonFly spinning-disk microscopy. WT-INF2 cells show a diffuse reticular network pattern, coinciding with the calreticulin distribution that generates a lace-like branching, polygon with a three-way junctions (TWJ) in the peripheral ER. The ER pattern is essentially similar to those untransfected cell (left side, asterisk). INF2 puncta often exist in vicinity of TWJ of peripheral tubular ER as well as perinuclear ER cisterna (asterisk). Mitochondria (gray) preferentially colocalize with the perinuclear ER cisterna. Bars = 10 µm and 2 µm.

**Figure 3.**
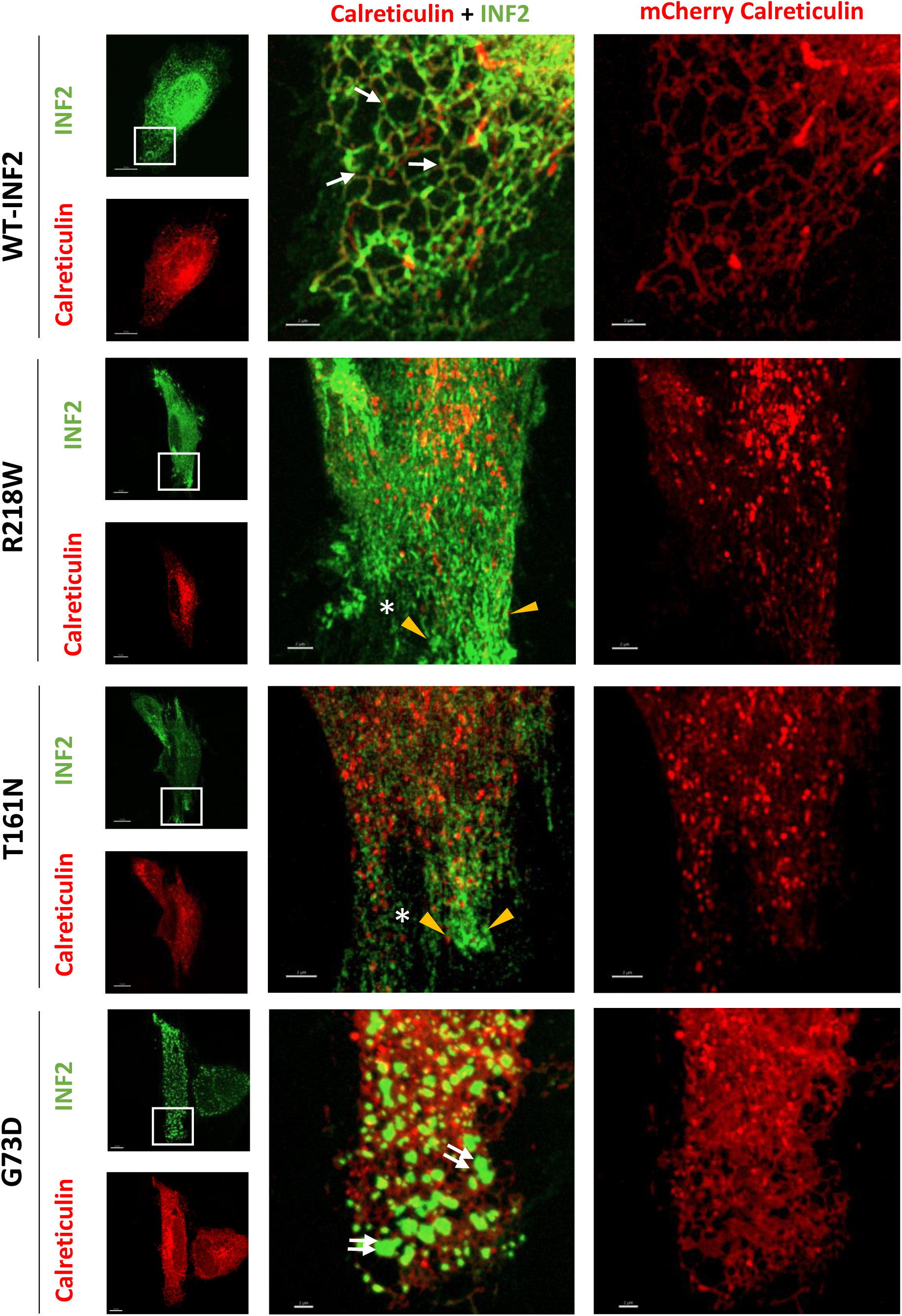
ER morphology in living HeLa cells expressing wild-type and pathogenic INF2 variants. GFP tagged, WT-INF2 (CAAX isoform) and its pathogenic variants (green) are transiently co-expressed with an ER-marker mCherry-Calreticulin (red) in HeLa cells. Cells expressing WT-INF2 show a typical tubular, peripheral ER pattern of a lace-like branching polygon with a three-way junction (white arrows). FSGS variants (R218W, T161N) strikingly alter ER tubular networks by introducing a focal compression (ER compaction, arrowheads), while leaving the polygonal structures intact in some area (asterisks). CMT+FSGS variant (G73D) cells show a diffuse ER fragmentation with tubular dilatation (double arrows) and sheet-like material accumulation. Bars = 10 µm and 2 µm.

### Actin organization in living HeLa cells expressing INF2 variants

To dissect the effect of the patient variants on the cytoskeletons, we then analyzed the actin network in living HeLa cells, which were double transfected with eGFP-tagged WT-INF2 or pathogenic variant (variant 1, CAAX ER-bound form), and an actin-marker Lifeact^12–14^. WT-INF2 cells produced abundant actin stress fibers both in the central region (central stress fibers) and along the cell border (peripheral stress fibers) (Fig 4). In contrast, FSGS variants (R218W, T161N) cells exhibited reduced the density and thickness of actin bundles, supplying fewer stress fibers in the cell center. CMT+FSGS variant (G73D) cells showed even fewer stress fibers compared to FSGS variant cells. The T161N variant demonstrated an intermediate cellular phenotype between FSGS and CMT+FSGS variants (Fig 4). Quantitative analysis revealed that the pathogenic variants reduce the stress fibers in the severity rank order: G73D > T161N > R218W and N202S (Sup Fig S4). Notably, we found the marginal actin bundles, which were more pronouncedly enriched in living INF2 variant cells labelled by LifeAct, might be quite transient, since they were invisible under the fixed condition and were devoid of bipolar vinculin-anchor (Sup Fig S5). We also observed a decrement in lamellipodia formation as well as cell migration activity in HeLa cells expressing INF2 variants (Sup Fig S6-8). Expression of depolymerization factor cofilin was not significantly altered in INF2 variant cells compared to the WT-INF2 cells (Sup Fig S15A). Moreover, INF2 pathogenic variant expressing cells exhibited longer filopodia, consistent with intrinsic biochemical property of INF2, which facilitates a linear-filament elongation^12,13^. Particularly, in CMT+FSGS variant cell, the filopodia were much longer but less mobile laterally than in FSGS variant cells (Sup Fig S9-10, Sup Mov S1). Aberrant INF2 CMT+FSGS variant punctate were found in antipodal cell protrusions including the shaft or tip of filopodia. Our data indicate that INF2 variants mainly dysregulate the linear actin filament formation implicated in remodeling of the cell cortex and protrusions, particularly at the cell poles (Sup Fig S20).

**Figure 4.**
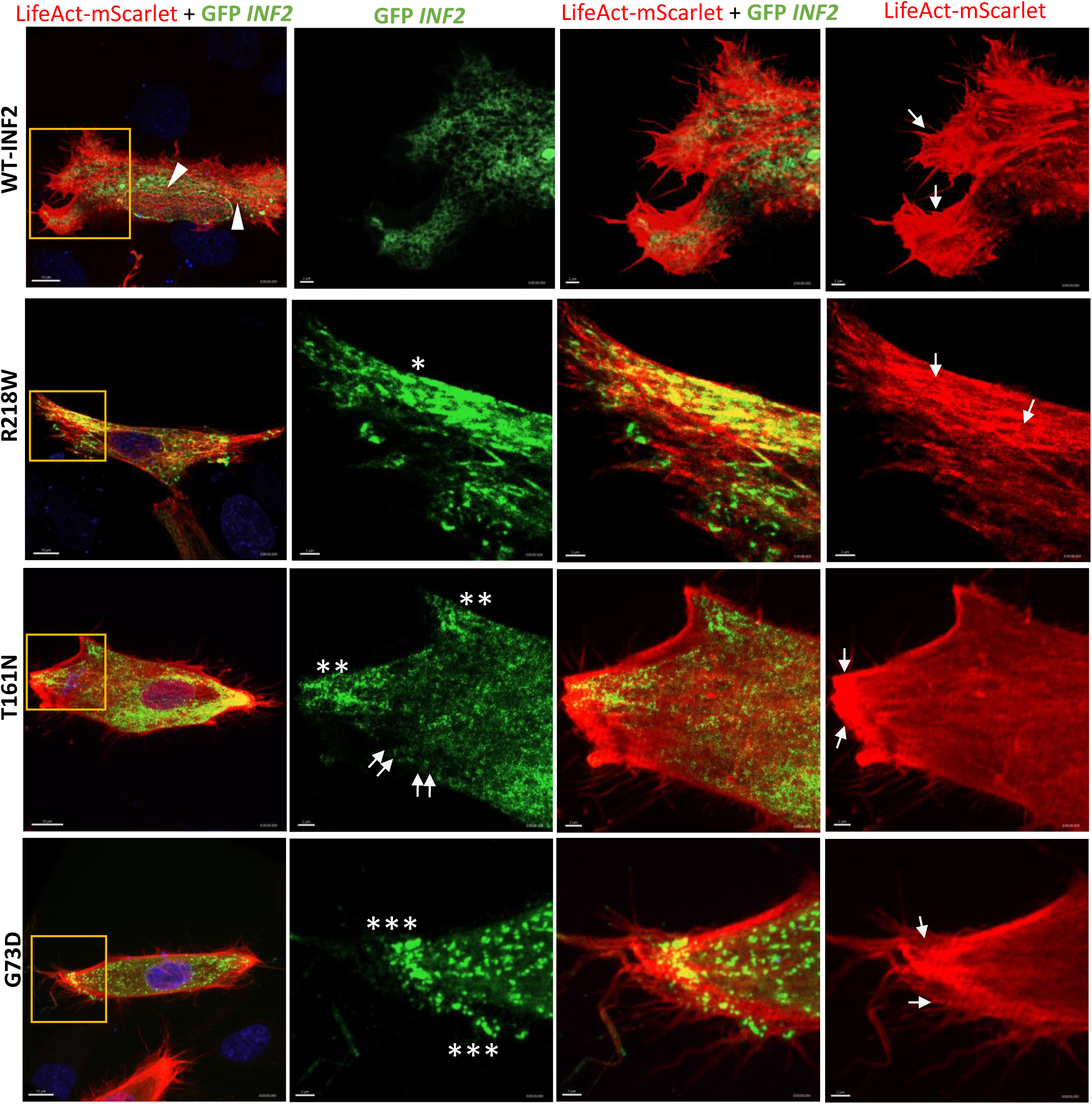
*INF2* and actin distribution in living HeLa cells expressing wild-type or pathogenic INF2 variants. GFP-tagged, WT-INF2 (CAAX isoform) and pathogenic variants (green) are transiently co-expressed with an actin-marker mScarlet-LifeAct (red) in HeLa cells. WT-INF2 cells display a diffuse reticular pattern with perinuclear enrichment. Actin fibers are abundant in the center (arrowheads) as well as along the cell edges (arrows). FSGS variant (R218W) cells display mild actin-disorganization, generating less central actin cables than WT-INF2 cells, while aberrantly accumulating the marginal bundles along the cell edges (arrows). Peripheral ER compaction (asterisks) are seen at the cell edges, coinciding with enrichment of peripheral actin bundles. In contrast, CMT+FSGS variant (G73D) cells produce shorter and thinner fibers, and accumulate coarse granular punctate in peripheral ER (triple asterisks). T161N variant cells show an intermediate actin phenotype between those of FSGS (R218W) and CMT+FSGS variant (G73D). There is mild accumulation of coarse matrices and focal ER compaction (double asterisks), while leaving some tubular ER network intact (double arrows). Bars = 10 µm and 2 µm.

We next examined the effects of the actin inhibitor (Cytochalasin D: CytoD) on peripheral tubular ER morphology using INF2 as a marker. CytoD shifted the ER balance towards a sheet-predominance in WT-INF2 cells, suggesting the roles of actin in shaping the tubular network under physiological conditions (Fig 5). In untreated T161N and G73D variant cells, the peripheral ER occasionally retracted from the cell margin, which might be due to un-tethering of ER from plasma membrane, and introduced a focal compaction (Fig 5). The CytoD treatment augmented the sheet/matrix accumulation as well as retraction in the peripheral ER of both T161N and G73D cells. Notably, CytoD treatment also affected the global cell shape in T161N, and G73D cells, but not WT-INF2 cells: CytoD partially relieved the cell lengthening of T161N and G73D cells, suggesting the elongation may reflect the dysregulated actin remodeling (Sup Fig S10, S12). Taken together, our observations revealed that INF2 variants reduce actin stress fibers (with focal adhesion), elongate the cell shape, and drive the ER-tubule-to-sheet transition than WT-INF2. The effects were more pronounced in CMT+FSGS than in FSGS variant-expressing cells.

**Figure 5.**
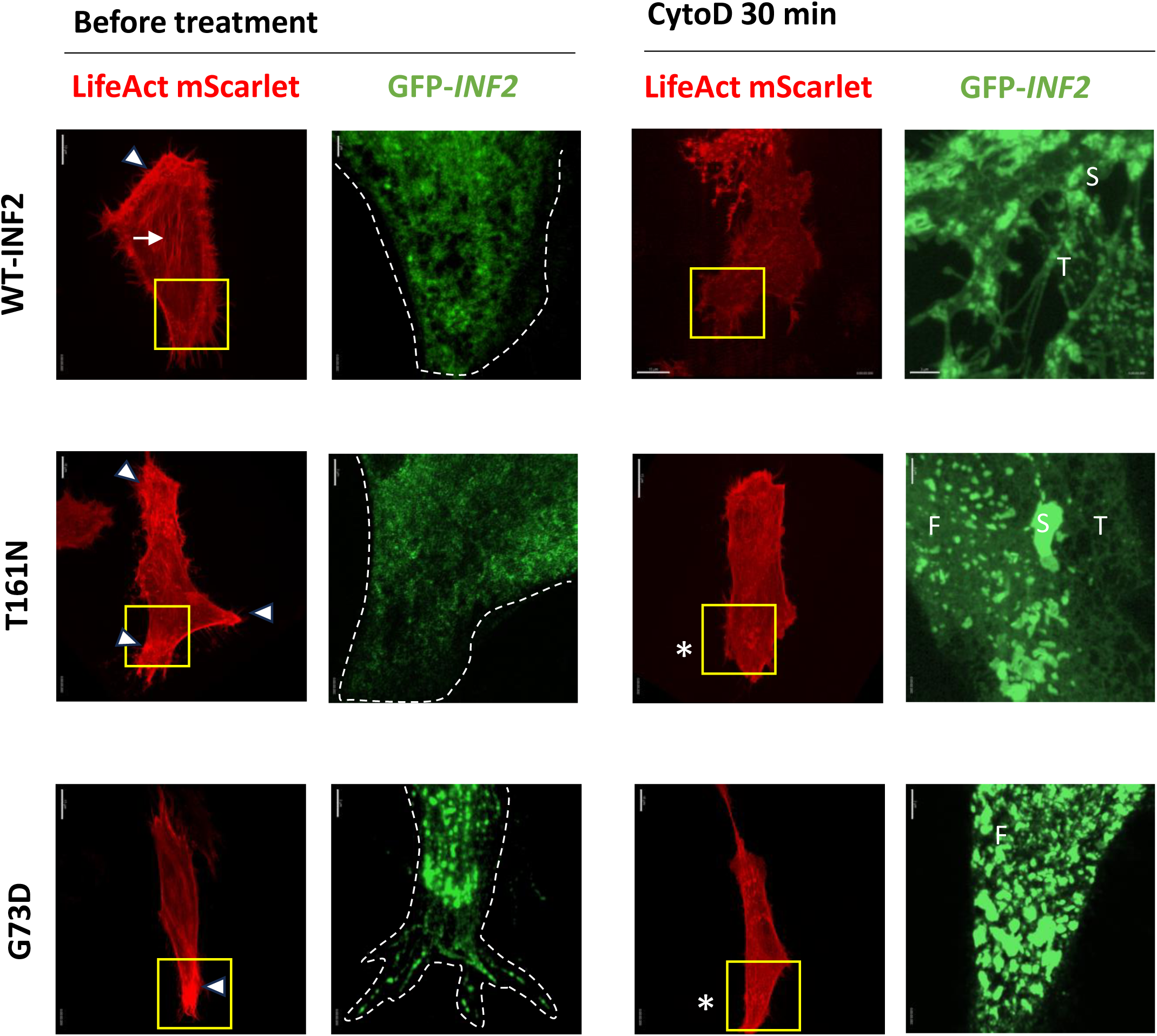
Effects of actin Inhibitior on the peripheral ER pattern in living HeLa cells expressing wild-type and pathogenic INF2 variants. HeLa cells are transiently transfected with eGFP-INF2 (WT, T161N and G73D) (green) and mScarlet-LifeAct (red). The effects of actin polymerization in the ER sheet-tubule balance are examined by treating cells with Cytochalasin D (CytoD) treatment (1µM for 30 min)^5^. Before CytoD treatment, cells expressing WT-INF2 generate a robust, central stress fibers (arrows) as well as peripheral bundles (arrowheads). T161N and G73D variant cells have elongated cell-shape with fewer and thinner central stress fibers. In contrast, peripheral actin bundles are focally enriched along the tip of cell protrusions often in an antipodal manner. The bundles remarkably attenuate in the presence of Cytochalasin D (asterisks). Boxed areas are magnified, and dot-lines indicate the contour of cell. Upon CytoD treatment, WT-INF2 cells alter the ER pattern as a sheet-predominant (S) over polygonal tubular structures (T). T161N cells preferentially increase a sheet-like ER component with some fragmentation (F). G73D cells generate a diffuse coarse-granular pattern and even more exaggerated fragmentation in response to CytoD. Bars = 10 µm & 3 µm.

### MT organization in living HeLa cells expressing INF2 variants

We next analyzed the MT arrays in living cells co-expressing INF2 variants and mScarlet-EMTB. WT-INF2 cells showed a radial MT arrays, which originated from the organizing center to the cell periphery and guided the ER strand to spread throughout the cytoplasm along the outer cell contour (Sup Fig S11). T161N and G73D variants tended to align the MT bundles in parallel to the long cell axis, thereby generating more peripheral ER retraction along the cell edge. In particular with G73D cells, ER tubules were often distended and fragmented, in addition to focal clustering of tubules, which were tightly surrounded by thick parallel MT bundles. Our data indicate that MT facilitates the ER shaping by anchoring them along the plasma membrane and spreading evenly in a lateral direction throughout the cell cortex (Fig 6), pathogenic INF2 variants impair the ER integrity through disorganization of MT arrays.

**Figure 6.**
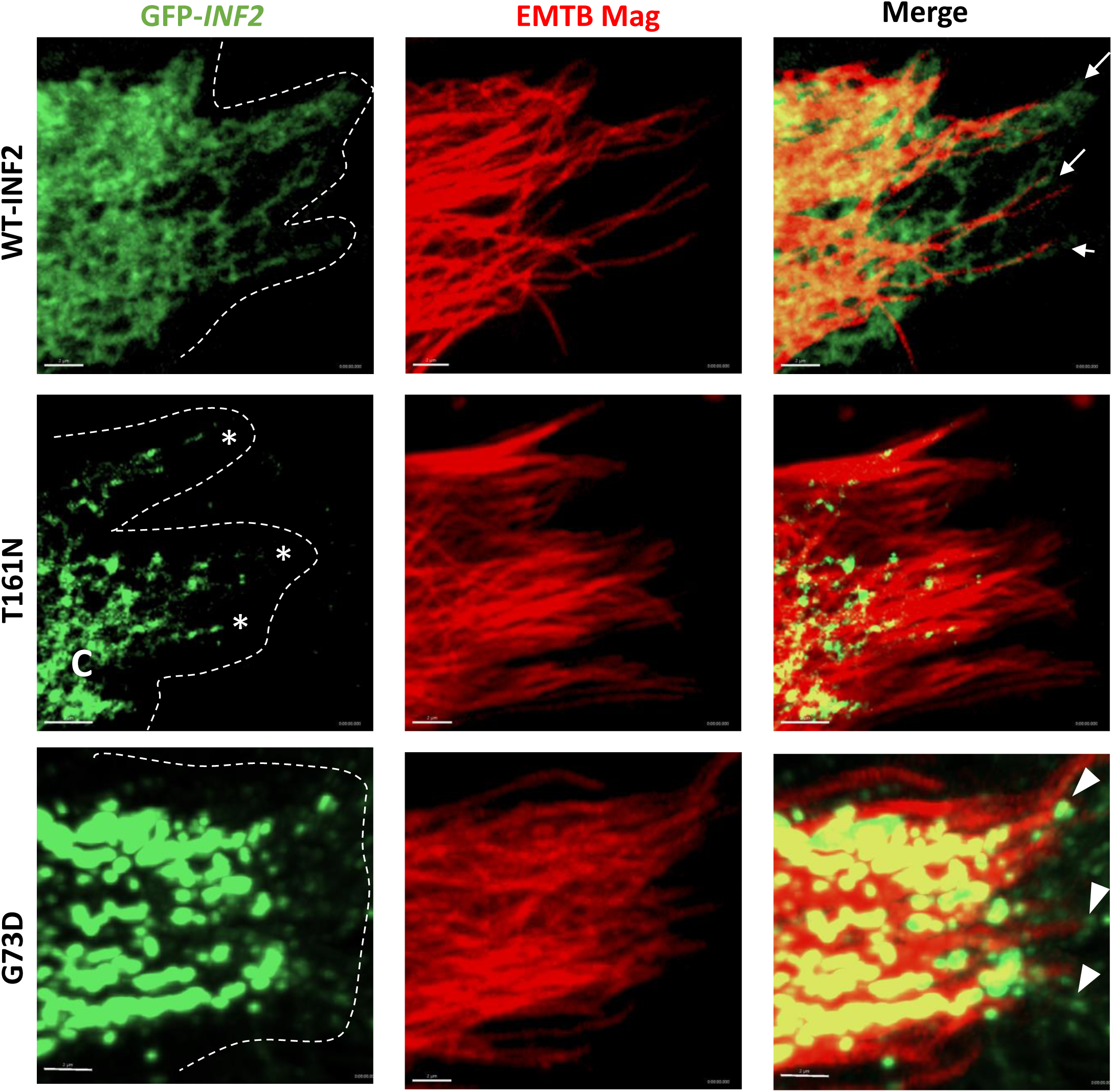
Interaction of ER with microtubule array in living HeLa cells expressing wild-type and *INF2* variants. Cells are co-transfected with eGFP-tagged WT-INF2, T161N, and G73D variants (green), and EMTB mScarlet (red). WT-INF2 cells showed a randomly projecting array of microtubules (MTs), which originate from the organizing center to periphery and guide the ER strand to expand along the plasma membrane (anchor point, arrows). Dot lines indicate the contour of the cell edge. T161N variant cells often align the MT bundles in parallel to the long cell axis, leading to the retraction ER from the cell edge (asterisks) as well as focal compaction (C) of polygons. In G73D variant cells, ER tubules are distended as well as fragmented. Aberrant G73D puncta are found in the cell poles including the shaft or tip of filopodia (arrowheads). The thick MT bundles align in parallel to the long axis, which compress the tubular networks. Bars = 2 µm.

Pathogenic INF2 variant cells altered microtubule arrays to align their bundle in parallel to the long cell axis, altering their cell shape to be more elongated “fusiform”. Treatment with a microtubule inhibitor, Nocodazole, did not significantly affect the cell length, but mildly broadened the cell width in both WT-INF2 and T161N, G73D variant cells (Fig 7, Sup Fig S12). In WT-INF2 cells, nocodazole mildly suppressed peripheral tubular formation as evidenced by the sparsity of polygonal meshwork. There was occasional enrichment of coarse-granular materials with sheet or matrix appearance, suggesting the need for MT to properly maintain the tubular polygon persistence. T161N and G73D cells had more frequent tubular ER compaction with peripheral ER retraction from the cell edge than WT-INF2 cells (Fig 7, Sup Fig S13). Nocodazole enhanced these tubule-to-sheet imbalance as well as retraction of ER. Notably, Nocodazole exerted more potent effects on the ER tubule-to-sheet transition than CytoD. The data indicate that the ER tubule-to-sheet balance primarily depends on interplay of ER with both actin and MT network. Pathogenic INF2 variants disrupted tubular network integrity through disruption of MT arrays, which maintain the tubule-to-sheet balance and expansion of peripheral ER.

**Figure 7.**
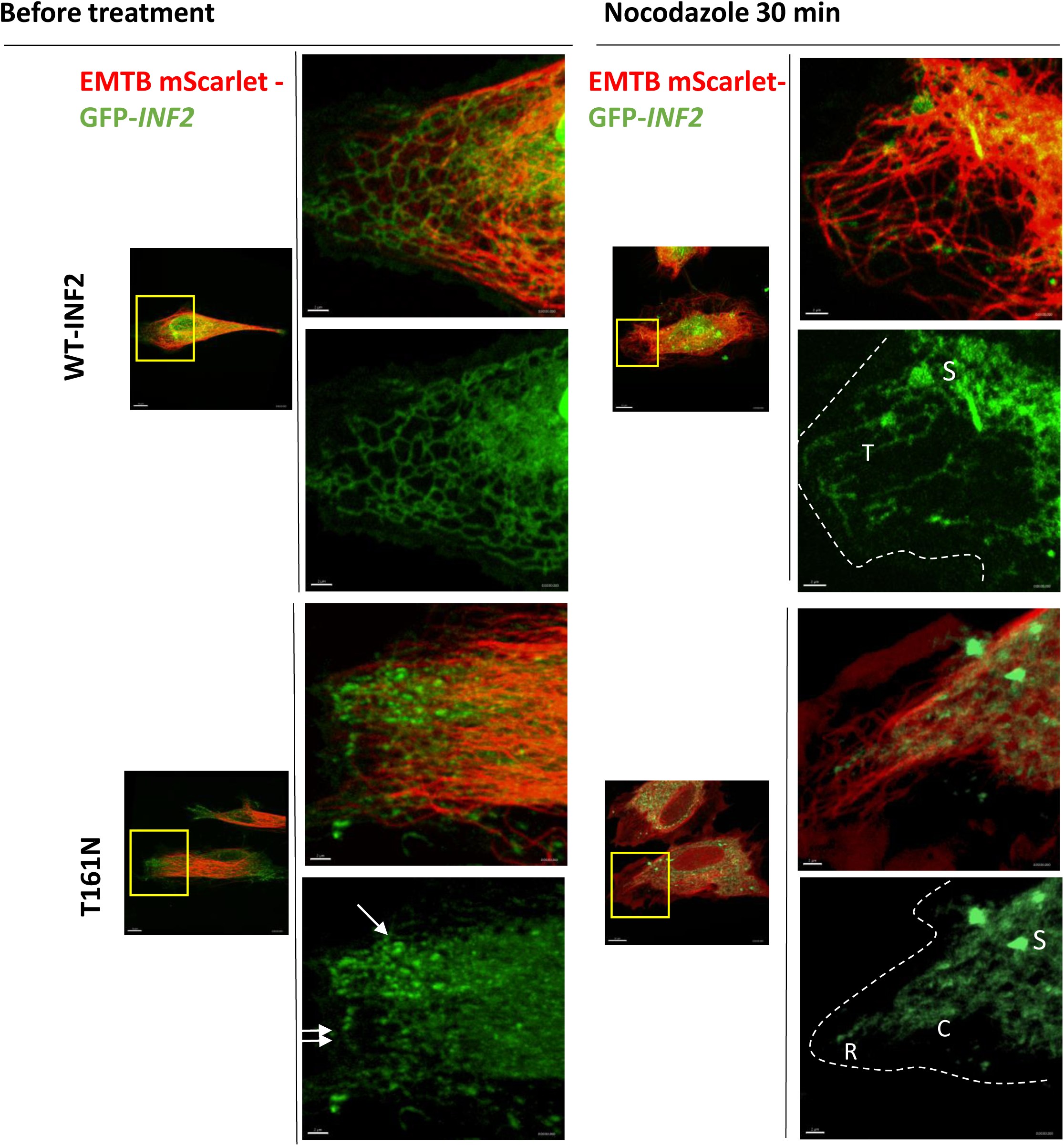
Effects of microtubule inhibitor on the peripheral ER in living HeLa cells expressing wild-type and pathogenic INF2 variants. HeLa cells are transiently cotransfected with eGFP-INF2 (WT, T161N) (green), and mScarlet EMTB (red). Cells expressing WT-INF2 show a reticular, tubular ER pattern along with a randomly projecting microtubule (MT) array. In contast, FSGS variant (T161N) cells often disarrange MT bundles parallelly aligned along the long cell axis, introducing focal compaction of tubule network (arrows), more frequently besides intact tubular network (double arrows). The effects of MT inhibitor in ER morphology are examined by treating cells with Nocodazole (Noc, 2.5 µg/ml) for 30 min. In WT-INF2 cells, Noc decreases peripheral tubular components (T) as evident by the sparsity of polygon meshwork, enrichment of sheet (S) or matrices materials. Upon Noc treatment, T161N cells have more frequent tubular ER compaction (C) with retraction (R) from the cell edge than WT-INF2 cells. Bars = 10 µm & 2 µm.

### Organelle-cytoskeleton interaction in living HeLa cells expressing INF2 variants

We next analyzed the mitochondria-ER interaction by labelling HeLa cells expressing eGFP-INF2 constructs and Mitotracker. In WT-INF2 expressing cells, the peripheral ER constituted a robust polygonal meshwork throughout the cell cortex^25,26^, which maintained the ER tubule continuity and the ER-mitochondria contacts with an appropriate distance (Fig 8). Mild FSGS variants (R218W and N202S) reduced peripheral ER polygons, which consequently diminished ER-mitochondria contact. G73D cells caused tubular dilation, fragmentation, and coarse-granular accumulation, indicating more potent degenerative effects on peripheral ER integrity and mitochondria interface (Fig 8A, Sup Mov S2). Our data indicate that INF2 variants perturbate the ER shape and distribution, leading to the reduction in effective contact sites. These effects were more pronounced in G73D than other variants with severing order of T161N > N202S, R218W.

**Figure 8.**
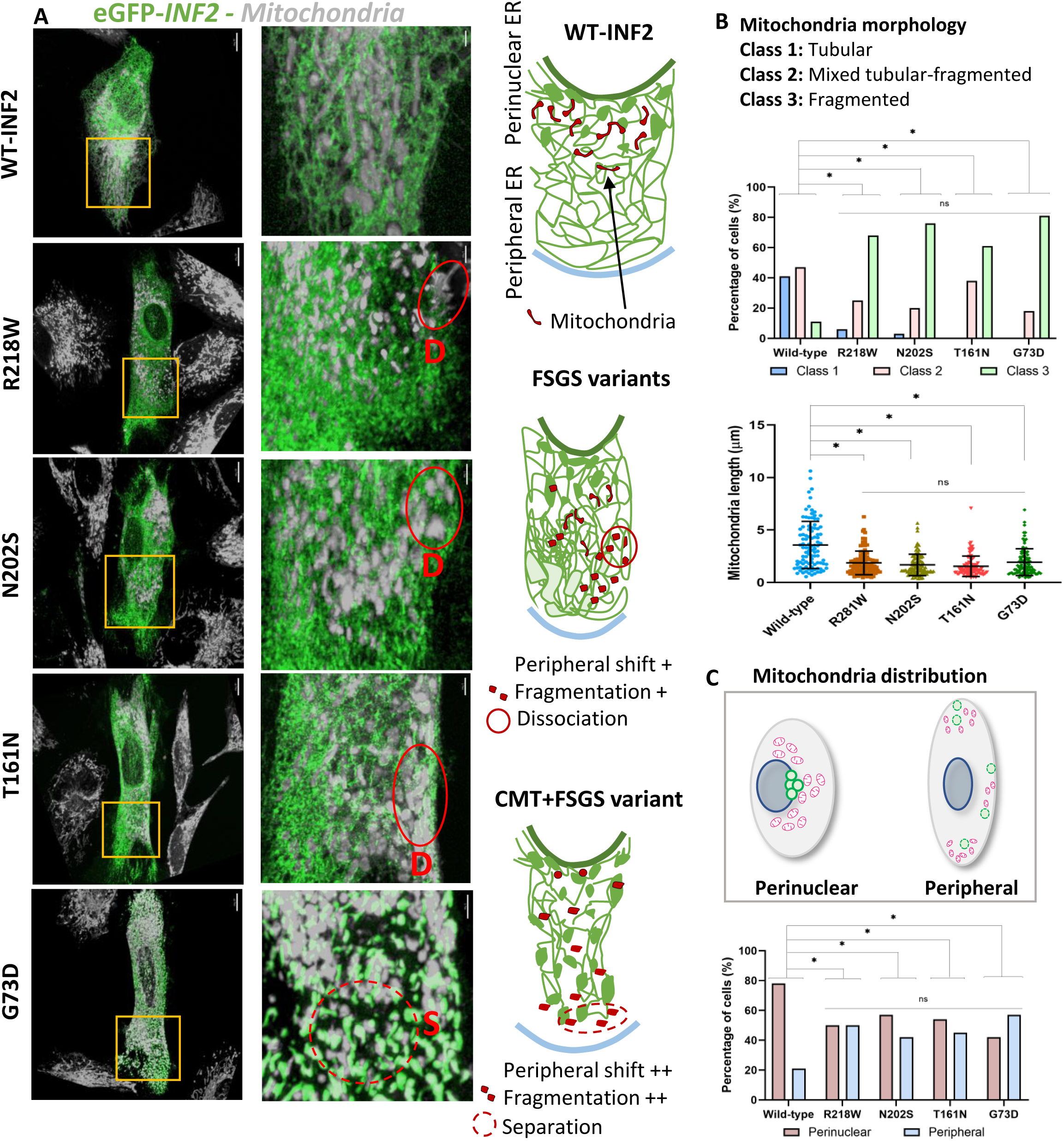
ER-mitochondria interaction in living HeLa cells expressing wild-type and pathogenic INF2 variants. **A. ER network visualized by eGFP-INF2.** Cell expressing WT-INF2 show a lace-like polygonal tubules in the peripheral ER, while those of FSGS variant cells (R218W, N202S and T161N) exhibited more frequent compaction. CMT+FSGS variant (G73D) cells show a ER fragmentation with tubular dilatation and sheet-like material accumulation. In WT-INF2 cells, mitochondria usually cluster in the perinuclear region and contact most closely with perinuclear ER cisterna. FSGS variants N202S, R218W cause a mitochondria shift to more periphery with fragmentation. T161N gives moderate deterioration inducing more frequent dissociation (D) of ER-mitochondria contact with fragmentation. G73D most severely impairs the contact of ER-Mitochondria, which exist separately (S). Bars = 10 µm and 2 µm. **B. Mitochondria shape:** Three classes of mitochondrial shape are: Class 1: predominant tubular-shape, Class 2: the mixture of tubular and fragmented mitochondria, and Class 3: fragmented. Mitochondria are more fragmented in INF2 variants (R218W, N202S, T161N, G73D) than those expressing WT-INF2. **(C) Distribution:** Mitochondria distribution pattern is subclassified into a binary category of either perinuclear (normal distribution) or peripheral (misdistribution). The data are from three independent experiments (n ≥ 30 cells per each variant).

Regarding the mitochondria shape, in WT-INF2 expressing cells predominantly showed a typical tubular-shape of mitochondria (Fig 8B). In contrast, pathogenic INF2 variant cells augmented mitochondrial fragmentation, particularly severe in CMT+FSGS variant cells compared to FSGS variants cells (T161N than N202S, R218W). Regards to the mitochondria distribution, mitochondria clustered predominantly in the perinuclear area in WT-INF2 cells, while misdistributing at the periphery more requently in INF2 variant cells (Fig 8C). In G73D cells, mitochondria more often clustered around cell periphery and occasionally retaining at the base of filopodia, the place where INF2 variants were concomitantly enriched (Fig 9). Our observations indicate that pathogenic INF2 variants alter mitochondria size, shape, and distribution in an actin-MT array dependent manner.

**Figure 9.**
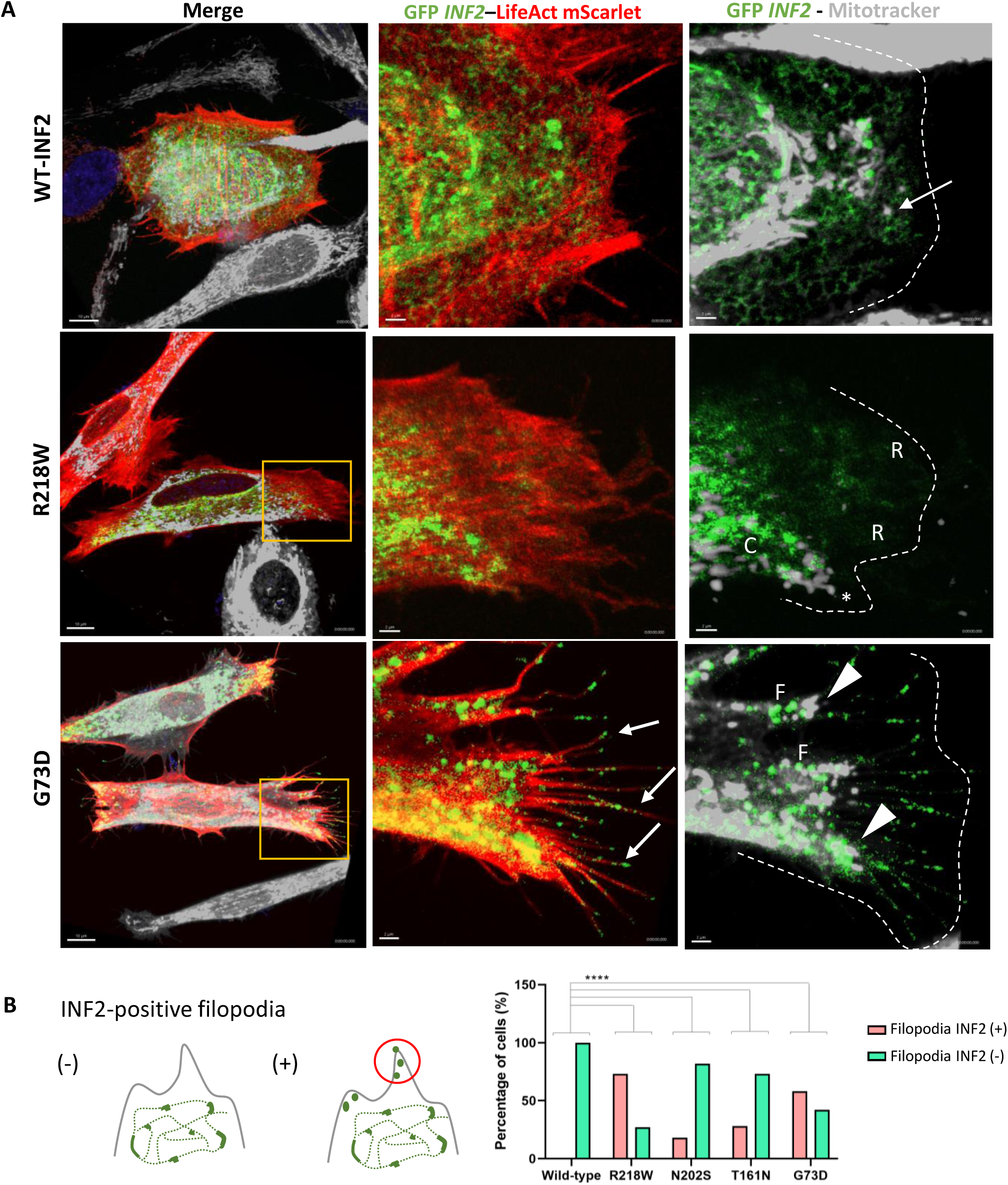
Mitochondria and INF2 localization in living HeLa cells expressing wild-type or pathogenic INF2 variants. **A, INF2 cells labeled with actin and mitochondria markers.** GFP-tagged, WT-INF2 (CAAX isoform) and pathogenic variants (green) are transiently co-expressed with an actin-marker mScarlet-LifeAct (red) and Mitotracker (gray). WT-INF2 cells cluster mitochondria along with the perinuclear ER cisterna (arrows). Intermediate variant (T161N) cells display a compaction (C) as well as retraction (R) of tubular ER and misdistribute mitochondria to the periphery (asterisk). In contrast, CMT+FSGS variant (G73D) cells show fragmented ER pattern (F), aberrant mitochondria retention at the base of the cell protrusion (arrowheads). G73D cells increase the number of filopodia. Note that enrichment of INF2 puncta in the tip or shaft of filopodia (arrows). Bars = 10 µm and 2 µm. **B. Frequency of the positive-INF2 staining in filopodia.** Representative filopodia pattern with INF2 puncta is illustrated. The frequency of filopodia with the positive INF2 puncta (filopodia lesion) is compared among cells expressing WT-INF2 and INF2 variants.

We next analyzed the effects of INF2 variant on the lysosome trafficking. Lysotracker labelling with HeLa cells expressing WT-INF2 revealed that there are two lysosome subpopulations: one is relatively immobile panel subpopulation constituting a perinuclear cluster around the MT organizing center (MTOC), and the other is a fast-moving lysosome at the periphery. Expression of intermediate (T161N) and the CMT+FSGS variant (G73D) in HeLa cells reduced lysosome traffic speed as well as total trajectory, compared with WT-INF2 cells (Sup Fig S14, Sup Mov S3). Trajectory analysis further revealed that lysosomal vesicles in G73D cells exhibited limited motility compared to the WT-INF2 cells. T161N cells showed the intermediate lysosome phenotype between WT-INF2 and G73D variants (Sup Fig S14). Our data indicate that INF2 variants impair the vesicle trafficking in the cell cortex, the degree of which correlated with the severity of cytoskeletal disarrangement.

### Functional mitochondrial deficits in HeLa cells expressing INF2 variants

To assess the effects of INF2 variants on mitochondria function, we measured oxygen consumption using the Seahorse flux analyzer. HeLa cells expressing WT-INF2 demonstrated the highest levels of basal respiration, maximal respiration, ATP production, and non-mitochondrial oxygen consumption. In contrast, the G73D cells exhibited the lowest values for all these indices including the lowest spare respiratory capacity, while T161N cells displayed the intermediate profile (Fig 10, Sup Fig S16-17). Our functional assays indicate that INF2 variants not only disrupt mitochondrial morphology and spatial distribution, but substantially perturbate the mitochondrial respiration.

**Figure 10.**
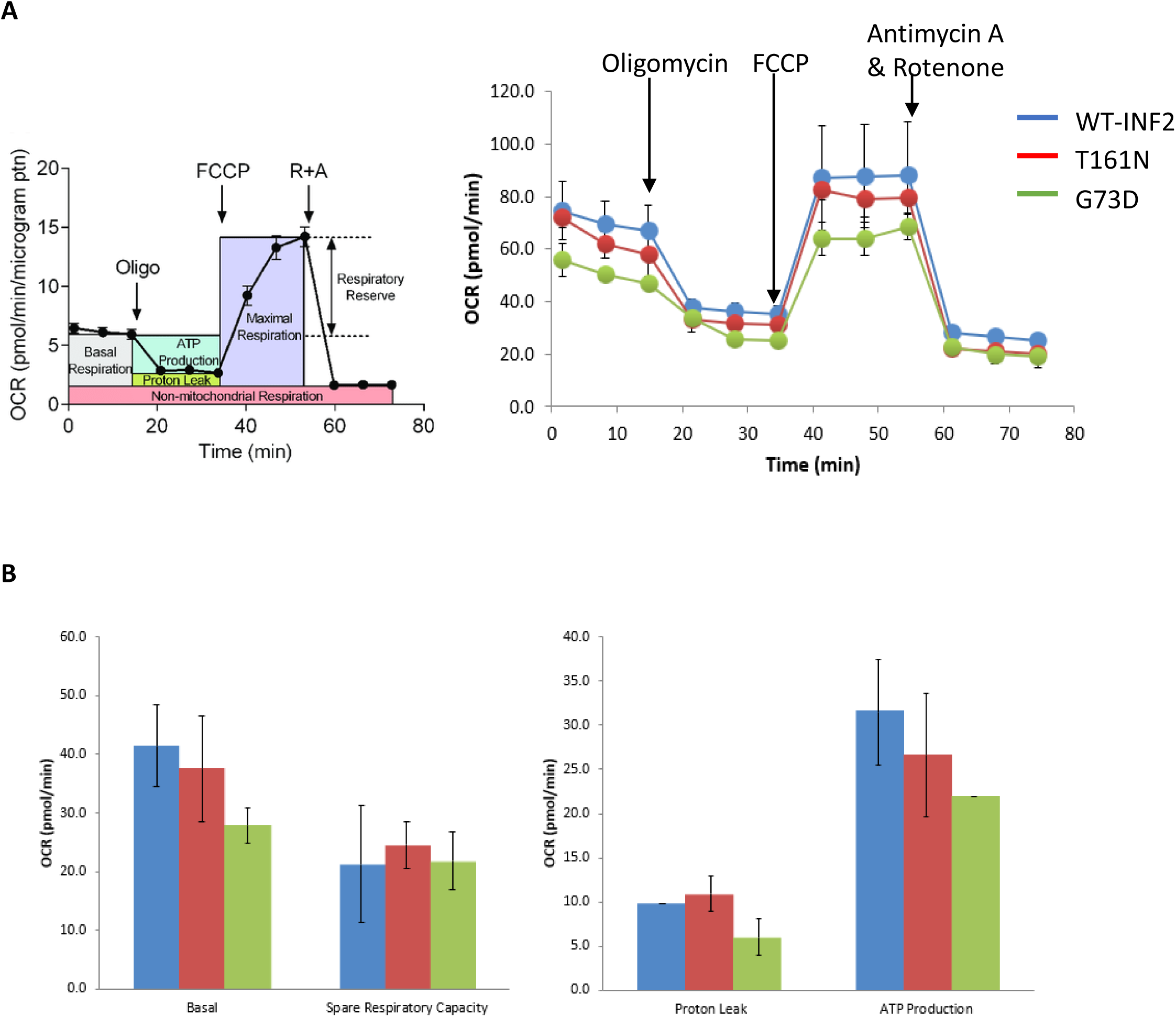
Impaired mitochondrial Respiratory Function in HeLa cells expressing *INF2* variants. HeLa cells are co-transfected with eGFP-tagged WT-INF2, T161N, and G73D variants. After 12 hours, cells are analyzed for mitochondrial function, by measuring their OCR in response to the three reagents. **(A) Diagram of Mitochondrial Respiration.** Cells are treated successively with complex inhibitor or coupling reagent (Oligomycin, FCCP and Rotenone + Antimycin A) in an automatically program in the Flux analyzer (XFp Agilent). Raw data are normalized and then analyzed by the WAVE software. **(B) Mitochondrial Respiration in *INF2* variants.** WT-INF2 cells exhibit a sound basal respiration as well as respiratory capacity during treatments, G73D cells show a decrease in the respiratory parameters, reflecting severely compromised mitochondrial function. T161N cells show the intermediate performance.

## DISCUSSION

Our present study, for the first time to our knowledge, revealed that INF2 pathogenic variants cause the ER disintegrity by using a high-resolution imaging of live cells. Pathogenic INF2 variants disintegrate dynamic tubular network, *e.g.* ER compaction and tubule-to-sheet transition. Particularly, CMT+FSGS variants induce more severe degeneration of ER including fragmentation and tubules dilation (Sup Fig S18). These ER disintegrity closely associates with the cytoskeletal disarrangement involving both actin and MT arrays, leading to the dysfunction of organelles like mitochondria and lysosomes (Sup Fig S19). The tubular ER network forms abundant membrane contact sites (MCSs) with other organelles (mitochondria, lysosome) and along the plasma membrane. The ER tethers the lysosomes and mitochondria to the membrane-bound compartment during the organelle traffic^4–6^, thereby regulating organelle dynamics, calcium homeostasis, and lipid metabolism. Mitochondrial respiratory deficits may play a central role in cell damage of both podocyte and Schwann cells, a hallmark phenotype in INF2 disorders.

### Role of INF2 mediated actin organization in ER integrity

ER disintegrity is a hallmark feature induced by the INF2 variants in our expression study with living HeLa cells. A tubule-to-sheet ER balance is maintained by coordinated interplay between ER and actin-MT network^1–6^. However, the actual mechanism for ER-actin interplay remains elucidated. Actin may act as a physical stabilizer at polygons or as-yet unknown factors could mediate the ER-actin interaction (Sup Fig S20)^5,27^. In our live-cell imaging, CMT+FSGS variant, which leads to the most severe disorganization of actin network, caused a drastic ER pattern change that converts tubules to sheet or matrix in the peripheral ER. FSGS variants R218W and N202S showed milder ER disintegrity, with focal tubular compression and sheet-like matrix enrichment at the place where the marginal actin bundles accumulate. Intermediate variant T161N led to moderate ER compaction, comparable in severity of concomitant actin disorganization. With regard to the tubular ER morphology, irreversible structural degeneration including diffuse fragmentation with dilatation of ER tubules were most prominent in G73D cells, followed by T161N, while N202S and R218W exhibited only milder clustering. The aberrant ER structural dynamics, such as the tubule-to-sheet ER transition, often accompany the concomitant cytoskeletal abnormalities, including cell elongation and accumulation of actin bundles at the cell margin. Our results taken together indicate that the INF2-DID variants impair the shaping and tubule-to-sheet balance of peripheral ER. Severe G73D variant further led to the substantial degenerative deficits, dilation and/or fragmentation.

### Effects of INF2 variants on actin network

The reduced actin stress fibers in our live cells expressing INF2 variants was essentially consistent with those with fixed HeLa cells or patient’s urinary epithelia previously reported^22,23^. INF2 variants (T161N, N202S and R218W) cells showed significantly less central actin cables (ventral fibers) than WT-INF2. Moreover, we observed fewer focal adhesions linking to thinner actin filaments with only atypical unipolar vinculin-anchorage at the cell edges (dorsal fibers), especially in G73D variant cells. Notably, the peripheral enrichment of actin bundle was relatively specific to our live cell expressing LifeAct and INF2 variants, and accumulated in an antipodal fashion for INF2 variant cells and thus might reflect their fusiform cell deformity. Such bundles were invisible under the fixed condition^23^. Therefore, it might due to immature and transient fibers, reflecting defective dynamic actin flow from the cell periphery to perinuclei (actin ring formation)^18,20,22,23^. Notably, pathogenic INF2 variant cells show the concomitant decrease in the cortical branched actin. Lamellipodia, another core actin-based structure, are formed by the branched actin networks via Arp 2/3 pathway^28^. We found that INF2 FSGS variants (T161N, N202S, R218W) mildly reduced the cytoplasmic punctate and discontinue the linearity of cortical actin than the CMT+FSGS variants, suggesting the role of INF2 in balancing the cortical actin network^29,30^. In concert with these defective actin assemblies and lamellipodia, cell migration was also impeded in cells expressing the pathogenic INF2 variant cells.

Elongation of the filopodia is one of the hallmarks of the pathogenic INF2 variants, which was only visualized by live imaging and might have been missed under fixation samples^21–23^. We found that cells expressing INF2 variants have elongated filopodia, at frequency order G73D > T161N > N202S, R218W. In WT-INF2 cells, stress fiber and filopodia formation was nearly abolished by CytoD, pointing the need of INF2 to activate actin assembly. A similar filopodia phenotype was reported for constitutively active mDia2, which accumulated long actin filaments and increased persistence of lamellipodial protrusion^31^. Moreover, G73D variant accumulated puncta in the tip or shaft of the elongated filopodia more frequent than intermediate (T161N) and mild FSGS variants (R218W, N202S). Ectopic INF2 expression in elongated filopodia may reflect defects in a series of linear actin-bundle formation starting from nucleation, elongation, to remodeling. These observations are in agreement with previous expression studies in HeLa and HepG2 cells^21,22^. Our data together with others indicate that INF2 variants dysregulate preferentially assembly of linear actin cables, while relatively suppressing the branched actin filaments (Sup Fig S21). The pathogenic effects are more pronounced for the CMT+FSGS variants than FSGS variants.

### Tissue-specificity of INF2 disorders

Podocytes exhibit a highly elaborated cell architecture with numerous foot processes covering the glomerular basement membrane, which are equipped with the dense actin network^11,30,32^. Regulation of the actin cytoskeleton is therefore critical for normal glomerular podocyte architecture and slit diaphragm function^30^. Schwann cells have also a unique structure consisting of myelinated multilayer sheath^11,30^. A fine-tuned, spatiotemporal remodelling of the actin cytoskeleton is necessary for proper myelin morphogenesis including radial sorting, wrapping and elongation of the Schwann cells along the axon^33^.

Our study supports the notion that the CMT+FSGS variants (residues 57 to 184 of DID) have more pronounced deleterious effects in cells than FSGS variants (residues 184 to 245 of DID). The unresolved question is how INF2 variants cause the diseases in two distinct cell lineages. Previous studies suggest that INF2 may not undergo canonical autoinhibition. The DID of INF2 self-interacts with the DAD competitively with actin monomer and binds each other in much weaker affinity than those of mDia (Kd 1.1 µM vs 0.28 µM)^34–36^. Notably, the naturally occurring pathogenic DID variants in the patients cause more prominent actin assembly defects than those of artificial variants of K792A polymerization, 3LA depolymerization-defective mutant^20^. This observation suggests that as-yet unknown INF2 interactors may participate in regulation of INF2 activity through heteromeric binding to the DID, in addition to the classic autoinhibition^20^. Taken together, the tissue selectivity of CMT phenotype may be due to the following possible INF2 interactions. First, INF2 may be regulated by proteins binding *in trans*^34^, Second, autoinhibition of INF2 might involve heteromeric proteins that facilitate the autoinhibition by promoting the DID-DAD interaction (*e.g.* CAP-KAC complex)^34^, Third, INF2 activity may vary depending on variable combination with tissue expression of formin isoforms (e.g., DAAM2)^37^. Fourth, INF2 activity may be altered by the balances between the local concentration of G-actin and its sequestration molecules and widely vary depending upon the cell type^16,35^.

### The role of INF2 in ER and actin-microtubule interaction

ER is the largest organelle in cells and contacts various organelles with physical continuity with the membrane-bound compartment^38^. ER shaping mechanisms, including atlastin-mediated stabilization of the peripheral tubular network, are vital for neuronal cells, which leads us to hypothesize that defective ER continuity might underlie the pathogenesis of disorders^39^. The prior studies pointed to the importance of MTs in ER spreading through anchoring peripheral ER tubules to the cell periphery^1–5^. MT arrays support the global ER extension over the cytoplasm and tether the peripheral ER to the plasma membrane. MTs closely associates with the tubular ER persistence^4^. MT plus ends drag the tubule ER along an existing MT shaft toward the growing cell leading edge (ER-sliding)^1,4,40^. INF2 directly colocalize and binds to MTs facilitating their bundling and stabilization^10,26,35,41^. Consistent with the previous reported INF2 properties, our MT inhibitor (Noc) study demonstrated that MT arrays are necessary for tubular persistence and tethering of the peripheral ER.

Actin dysregulation also plays a critical role in the loss of ER integrity caused by INF2 variants. First, actin disorganization disrupts the MT array by perturbating the efficient MT capture to cortical actin. Second, we found a focal ER compaction coincide with the peripheral enrichment of actin bundles. Third, actin network may affect ER pattern by lengthening the global cell shape with the concomitant MT parallel alignment. We found that cells expressing INF2 variants align MTs in parallel to the long cell axis, resembling HeLa cells expressing constitutively active mDia1 mutant (ΔN3)^42^. CytoD relieved the cell elongation of INF2 variants (T161N, G73D) but not Noc. The parallel alignment of MT might thus contribute to the compaction of ER secondarily to the defective actin network. Fourth, actin flows simultaneously draw ER back toward the nucleus and stabilize the perinuclear ER^1^. Some ER extension operates solely depending actin filament (budding extension)^1^. Treatment with Nocodazole also cause milder ER changes than CytoD. Taken together, our data suggest that actin mainly regulates dynamic tubule-to-sheet balance, as well as the global extension pattern of ER in co-operation with MT.

### Defective mitochondria, functions and lysosome trafficking in INF2 disorder

Mitochondrial dysfunction significantly impacts on the pathogenesis of various human renal and neuronal disorders^6,43^. In cells expressing pathogenic INF2 variants, mitochondria misdistribute to the cell periphery, suggesting the importance of coordinated MT and actin organization in positioning mitochondria properly around the perinuclear area^44^. The aberrant mitochondria pattern is more pronounced in cells expressing CMT+FSGS variants compared to FSGS variants, which correlates with the severity of cytoskeletal disorganization as well as ER disintegration. Our observations indicate that dysregulated actin-MT networks and ER-organelle contacts likely closely link to these mitochondrial defects.

Mitochondrial activity relies upon a proper fission and fusion cycles that maintains mitochondrial homeostasis^43,45^ Mitochondrial fission at ER-mitochondria contact sites relies on the local actin filament assembly. An ER-resident, INF2-CAAX isoform initiates a fission event independently of Drp1 and enhances the ER-to-mitochondria Ca transfer^15^. Pathogenic INF2 variants affect the ER-mitochondria contacts, interfering these two-organelle interface and/or local actin dynamics crucial for the optimal fission^6,14^. Our analysis demonstrated that G73D and T161N variants decrease the basal and maximal mitochondrial respiration, ATP production, and proton leak. Podocytes and peripheral nerves, with a high energy demand, depend on proper mitochondrial dynamics. Mutations affecting mitochondrial fission, such as MFN2, OPA1, and GDAP1, significantly contribute to the three subtypes CMT^46,47^. Taken together mitochondria dysfunction could play a key role in cell damage in INF2 disorders.

Actin filaments serve as a scaffold network for cellular cargo transport in the cell cortex. Our trajectory analysis revealed significantly limited lysosome motility in T161N and G73D cells compared to WT-INF2 cells. Peripheral lysosomes in the cell cortex undergo a long-range vesicle trafficking via MTs using kinesin and dynein motors, while short-range, fast transport occurs on actin filaments by myosin motors^48,49^. INF2 is implicated in actin comet formation for the trafficking vesicles in the cell cortex. Moreover, MT facilitates organelle transportation in co-operation manner with motor proteins or ER sliding mechanism^21,24^, and thus maintains the homeostasis of slit-diaphragm complex trafficking^50^. INF2 binds to MAL2, the complex of which are necessary for apical transcytosis as well as lateral lumen formation in hepatoma HepG2 cells^21^. Pathogenic INF2 variants disrupt their heterometric interaction with dynein light chain 1, misdirecting dynein-mediated post-endocytic sorting of nephrin^50^. Our data, together with others, indicate that lysosome motility is perturbated by INF2 variants, thereby comprising one of the core mechanisms of cellular dysfunction in INF2 disorder.

## Conclusion

Our live cell imaging revealed that the pathogenic INF2 variants induce aberrant ER morphology and distribution in an actin-MT dependent manner (Sup Fig S22-23, Table 1). Such ER disintegrity leads to organelle dysfunction through disruption of ER-organelle contacts, thereby diminishing mitochondria respiration and lysosome trafficking, which ultimately disable the cell viability. Further study is necessary to clarify the role of tissue-specific factors and underlying autoregulatory mechanisms.

## SUPPLEMENTARY METHODS

### Locations of INF2 variants in single FSGS and CMT+FSGS dual disease

The distribution of variant positions was consistent with the previously reported genetic landscape: all the human *INF2* variants exclusively clustered in the DID domain, the proximal DID variants (acid residue 57 to 184) cause CMT+FSGS, while the distal ones (residue 184 to 245) give a rise to a single FSGS phenotype^11^. Among these, we chose four INF2 variants from three subclasses: G73D for CMT+FSGS, T161N for the intermediate, and N202S and R218W for mild FSGS (Fig 1A). T161N variant is located in the boundary region, which could cause both CMT+FSGS and FSGS phenotypes^11^. Segregation of two clinical phenotypes (dual CMT+FSGS or single FSGS) into two separated regions (proximal vs distal, respectively) suggests that the N-terminal portion of DID may mediate a specialized interactions with other heteromeric binding partners in Schwann cell.

### Live cell imaging by the spinning disk confocal microscopy

HeLa cells were plated at a density of 3x10^4^ per 35 mm glass-bottom dish and were transfected with N-eGFP-INF2 cDNA3.1 with the cytoskeleton-marker plasmids (either EMTB or LifeAct) using TransIT-LT1 (MIR 2300, Mirus) accordingly the manufacturer’s instruction. Eight to ten hours after the transfection, cells were processed for immunofluorescent staining, western blotting, or flux analyzer.

INF2 variant-expressing cells were visualized using a high-resolution spinning disk microscopy (DragonFly) with a 100X lens, capturing a total time-lapse of 10-20 seconds with 1-second increments. Laser intensity and exposure time were optimized for each experimental conditions to minimize photobleaching toxicity. Images were then subjected to a deconvolution process, which enabled the image to gain more sharpness and higher resolution.

### Markers for cytoskeleton and organelles

ER was visualized by mCherry-Calreticulin (Addgene, #55006). To label the actin filaments in live cells, LifeAct-mCherry was obtained from Addgene (#193300). LifeAct-mScarlet-I was generated by replacing the mCherry with mScarlet, which derived from the ITPKA-mScarlet-I (Addgene #98829). For the microtubule live imaging, microtubule-binding domain of ensconsin (EMTB)-mScarlet-I construct was constructed by replacing the 3xGFP of EMTB-3xGFP (Addgene #26741) with mScarlet-I.

To visualize the mitochondria in living cells, cells were labelled with either MitoTracker Red CMXRos (M7512, Invitrogen) or MitoTracker Deep Red FM (M22426, Invitrogen) in a final concentration of 50 nM for 30 minutes. Cells were marked for lysosomes by LysoTracker Red DND-99 (L7528, Invitrogen) with 50 nM dilution and were thereafter kept under the complete medium during imaging procedure. Cell nuclei were stained with Hoechst 33342 (346-07951, Dojindo).

### Antibodies

Anti-Cofilin (D3F9) is a rabbit monoclonal antibody against peptide corresponding to central residues of human cofilin1 protein (#5175, Cell Signaling) (Sup Fig S13). Anti-GADPH is a mouse monoclonal antibody against Glyceraldehyde-3-phosphate dehydrogenase (014-25524, Fujifilm). Anti-cortactin [EP1922Y] is a rabbit monoclonal antibody raised against Cortactin (ab81208, Abcam). Phalloidin-iFluor 647 Conjugate is from Cayman Chemical Co. (Item No. 20555). For the secondary antibody in immunocytochemistry, Donkey anti-Rabbit IgG was obtained from Invitrogen (#A-31572, 1:1000). For the secondary antibody in western blot, Anti-Mouse IgG, HRP-Linked Whole Ab Sheep (Cytiva, NA931) and Anti-Rabbit IgG, HRP-Linked Whole Ab Donkey (Cytiva, NA934) were used.

### Morphometry of Cell shape

The length of typical spindle-shaped HeLa cells was defined by measuring the longest longitudinal distance between two points at the cell edges. In cells with more round shape, the length is defined as the greatest distance between any two points along the cell margin (Feret’s diameter).

### Immunofluorescence staining for cortactin and cofilin

Expression of cortactin and cofilin in INF2 variants expressing cells was studied by indirect immunofluorescence in HeLa or COS-7 cells after fixation with 4% paraformaldehyde. Cells were then blocked in 5% goat serum in PBS and incubated successively with the primary antibodies of rabbit anti-cofilin (dilution 1:500) or cortactin (dilution 1:1000), followed by the secondary antibody of Alexa Fluor 555 Donkey anti-Rabbit IgG (dilution 1:1000). After mounting, the samples were observed under a high-resolution, spinning-disk microscope DragonFly.

Following the quantification method described in elsewhere^29^, after the immunofluorescence staining, images were captured with a spinning disk confocal microscope. Images of INF2 variants were imported and analyzed by the Fiji program. Immunostaining of cortactin in COS-7 cells was used for the quantification. Background fluorescence was subtracted from the image, and individual cells were then digitally outlined with Fiji software (National Institutes of Health) to measure total cell fluorescence for cortactin. To quantitate the fraction of the lamellipodia area, the occupancy of the cortactin-positive regions and actin-rich fringes was measured^50^ in total cytoplasmic area, yielding the relative area of cortical actin meshwork. A fraction of the selected lamellipodia regions in the cortactin-positive cell cortex areas were expressed as a percentage occupancy in the total cell area. To obtain the relative length of the cortactin-positive membrane, we measured the length of the plasma membrane enriched with cortactin by digitally outlining the rim cortactin signal along the leading edge. Then, a percentage of the summed positive cortactin length per total cell perimeter was calculated. The both relative area of cortical actin mesh and the relative length of the cortactin-positive membrane were examined for at least 20 cells for each variant.

### Western Blot analysis

Ten hours after transfection with eGFP-INF2 (wild-type or pathogenic variants T161N and G73D), HeLa cells were trypsinized and homogenized in cold homogenization RIPA buffer (182-02451, Watson) supplemented with complete protease inhibitor cocktail (163-26061, Watson) in 30 minutes. Cell lysates were clarified by centrifugation at 800 *g* for 5 minutes at 4°C. Supernatants were collected and the protein concentrations were determined by using Bio-Rad DC Protein Assay. Equal amounts of protein (10 µg) were loaded on Bolt 4-12% Bis-Tris Plus Gel (NW04120, Invitrogen) and separated by electrophoresis. The proteins were transferred to the PVDF membrane (Amersham) by use of a semi-dry Trans-Blot SD cell (BioRad). Cofilin expression was analyzed by use of anti-Cofilin antibody (Rabbit, dilution 1:1000). GADPH was used for an internal control (Mouse anti-GADPH, dilution 1:1000). Protein band intensities were measured using ImageJ software and calibrated by internal control of GADPH.

### Cell migration assay

A confluent monolayer of HeLa cells was seeded at 3x10^4^ cells in a 35mm dish). After transfection 10 hours, cells were mechanically wounded (‘scratch assay’) with a sterile pipette tip, leaving two wound edges (dashed lines) separated by a void. The Cell migration is qualitatively assessed by visual inspection. Cells at the leading edges quickly adopted a polarized morphology and formed broad lamellae pointing into the direction of the void are judged as “migrating phenotypes”. Otherwise, cells were annotated as stationary (white dash-line circle), or vanishing phenotype (appears only within two time-frame).

## ACKNOWLEDGEMENTS

The authors are grateful to all the patients, their family members, and reference physicians, who participated in our study. We thank Dr. Shinji Hirano (Department of Cell Biology, Kansai Medical University) for providing antibodies against the focal adhesion and Dr. Ryu Takeya (Department of Pharmacology, Falculty of Medicine, University of Miyazaki) for helpful discussion. This work was supported by the Grants-in-Aid for Scientific Research (C) (23K07730 to H. T) from the Japan Society for the Promotion of Science, and the Uehara Memorial Foundation Research Fellowship Program and KMU Mizuno Takako award to TTHQ.

## AUTHOR CONTRIBUTION

HT conceived and designed this study. QTHT and HT wrote the manuscript. QTHT, NK, and HU performed cell biological studies. YM supported the mitochondrial function analysis. All authors read and approved the final manuscript.

## DATA SHARING STATEMENT

The datasets generated and/or analyzed during the current study are available in the temporary review repository at https://drive.google.com/drive/folders/15NsMyMrvPFAAUmTjN8KH7v6E1sAIkPEX

---

**Supplementary Table S1.** Summary of effects of INF2 mutations on ER integrity in Live HeLa Cells.

| INF2 variants<br>Pathogenic severity | Wild-type | R218W, N202S<br>Mild | T161N<br>Moderate | G73D<br>Severe |
| --- | --- | --- | --- | --- |
| Peripheral ER |  |  |  |  |
| Tubule-sheet dynamics | Tubule | Tubule > sheet | Tubule > sheet | Sheet > tubule |
| Spatial arrangement | Perinuclear | Focal compaction in peripheral ER | Global compaction (+) in peripheral ER | Global compaction (++) in peripheral ER |
| Tubular morphology | • Polygons with TWJ (+++) | • Polygon (++)<br>• Conversion of tubule to matrix or sheet (+) | • Polygon +<br>• Conversion of tubule to matrix or sheet (++)<br>• Dilatation with fragmentation (+) | • Dilatation with fragmentation (++) |
| Cytoskeleton |  |  |  |  |
| Actin |  |  |  |  |
| Central stress fiber | (+++) | (++) | (++) | (+/-) |
| Peripheral bundle along cell edge | (±)<sup>a</sup> | (+) | (+) | (+) |
| Effects of Cytochalasin D | ER Sheet > Tubule | ND | ER Sheet > Tubule | No morphologic change<sup>b</sup> |
| Microtubule | • Radial arrays<br>• Apparent MTOC |  | • Parallel > Radial arrays<br>• Ambiguous or loss of MTOC | • Exclusively parallel bundle<br>• Loss of MTOC |
| Effects of Nocodazole | • ER retraction<br>• Focal spareress in ER polygons<sup>c</sup> | ND | • ER retraction<br>• No morphologic changes | • ER retraction<br>• No morphologic changes |

(a) WT-INF2 cells generate some marginal fibers in amount less than pathogenic INF2 variant cells. The exaggerated marginal fibers in the pathogenic variant cells may be due to focal accumulation of highly dynamic actin filaments, because they are invisible under the fixed conditions.
(b) The predominant feature of tubular fragmentation seems to represent a substantial degeneration of tubular structure, because it does not restore their morphology in response to CD treatment
(c) Partial recovery attained after withdrawal of Nocodazole, suggesting the reversibility.
MTOC: Microtubule organizing center, ND: not determined

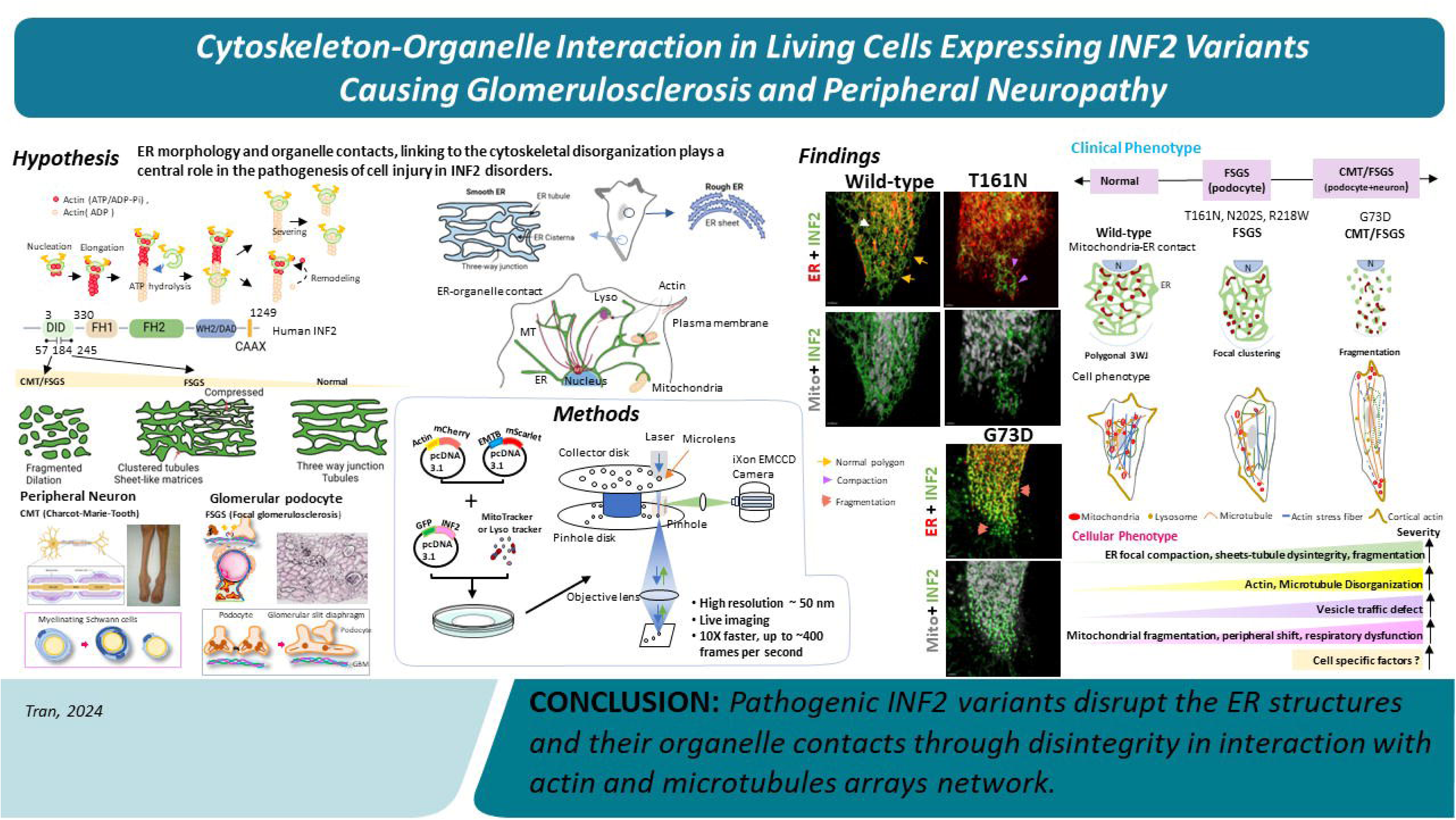

## Notes

### Competing Interest Statement

The authors have declared no competing interest.

## REFERENCES

1. Obara CJ, Moore AS, Lippincott-Schwartz J. Structural diversity within the endoplasmic reticulum—from the microscale to the nanoscale. CSH PERSPECT BIOL. 2023;15(6):a041259.

2. Shibata Y, Shemesh T, Prinz WA, Palazzo AF, Kozlov MM, Rapoport TA. Mechanisms determining the morphology of the peripheral ER. Cell. 2010;143(5):774–788.

3. Zheng P, Obara CJ, Szczesna E, et al. ER proteins decipher the tubulin code to regulate organelle distribution. Nature. 2022;601(7891):132-138.

4. Wu H, Carvalho P, Voeltz GK. Here, there, and everywhere: The importance of ER membrane contact sites. Science. 2018;361(6401):eaan5835.

5. Joensuu M, Jokitalo E. ER sheet–tubule balance is regulated by an array of actin filaments and microtubules. Exp Cell Res. 2015;337(2):170–178.

6. Manor U, Bartholomew S, Golani G, et al. A mitochondria-anchored isoform of the actin-nucleating spire protein regulates mitochondrial division. eLife. 2015;4:e08828.

7. Rottner K, Faix J, Bogdan S, Linder S, Kerkhoff E. Actin assembly mechanisms at a glance. J Cell Sci. 2017;130(20):3427–3435.

8. Wallar BJ, Alberts AS. The formins: active scaffolds that remodel the cytoskeleton. Trends Cell Biol. 2003;13(8):435–446.

9. Chhabra ES, Higgs HN. The many faces of actin: matching assembly factors with cellular structures. Nat Cell Biol. 2007;9(10):1110–1121.

10. Hegsted A, Yingling CV, Pruyne D. Inverted formins: A subfamily of atypical formins. Cytoskeleton. 2017;74(11):405–419.

11. Labat-de-Hoz L, Alonso MA. The formin INF2 in disease: progress from 10 years of research. Cell Mol Life Sci. 2020;77(22):4581–4600.

12. Chhabra ES, Higgs HN. INF2 Is a WASP homology 2 motif-containing formin that severs actin filaments and accelerates both polymerization and depolymerization. J Biol Chem. 2006;281(36):26754–26767.

13. Ramabhadran V, Gurel PS, Higgs HN. Mutations to the formin homology 2 domain of INF2 protein have unexpected effects on actin polymerization and severing. J Biol Chem. 2012;287(41):34234–34245.

14. Korobova F, Ramabhadran V, Higgs HN. An actin-dependent step in mitochondrial fission mediated by the ER-associated formin INF2. Science. 2013;339(6118):464-467.

15. Chakrabarti R, Ji W-K, Stan RV, de Juan Sanz J, Ryan TA, Higgs HN. INF2-mediated actin polymerization at the ER stimulates mitochondrial calcium uptake, inner membrane constriction, and division. J Cell Biol. 2018;217(1):251–268.

16. Goode BL, Drubin DG, Barnes G. Functional cooperation between the microtubule and actin cytoskeletons. Curr Opin Cell Biol. 2000;12(1):63–71.

17. Rodriguez OC, Schaefer AW, Mandato CA, Forscher P, Bement WM, Waterman-Storer CM. Conserved microtubule–actin interactions in cell movement and morphogenesis. Nat Cell Biol. 2003;5(7):599–609.

18. Brown EJ, Schlöndorff JS, Becker DJ, et al. Mutations in the formin gene INF2 cause focal segmental glomerulosclerosis. Nat Genet. 2010;42(1):72–76.

19. D’Agati VD, Kaskel FJ, Falk RJ. Focal segmental glomerulosclerosis. NEJM. 2011;365(25):2398–2411.

20. Boyer O, Nevo F, Plaisier E, et al. INF2 mutations in Charcot–Marie–Tooth disease with glomerulopathy. NEJM. 2011;365(25):2377–2388.

21. Madrid R, Aranda JF, Rodríguez-Fraticelli AE, et al. The formin INF2 regulates basolateral-to-apical transcytosis and lumen formation in association with Cdc42 and MAL2. Dev Cell. 2010;18(5):814–827.

22. Bayraktar S, Nehrig J, Menis E, et al. A deregulated stress response underlies distinct INF2-associated disease profiles. JASN. 2020;31(6):1296–1313.

23. Ueda H, Tran QTH, Tran LNT, et al. Characterization of cytoskeletal and structural effects of INF2 variants causing glomerulopathy and neuropathy. Sci Rep. 2023;13(1):12003.

24. Tikhomirova MS, Kadosh A, Saukko-Paavola AJ, Shemesh T, Klemm RW. A role for endoplasmic reticulum dynamics in the cellular distribution of microtubules. PNAS. 2022;119(15):e2104309119.

25. Ramabhadran V, Korobova F, Rahme GJ, Higgs HN. Splice variant–specific cellular function of the formin INF2 in maintenance of Golgi architecture. Mol Biol Cell. 2011;22(24):4822–4833.

26. Gaillard J, Ramabhadran V, Neumanne E, et al. Differential interactions of the formins INF2, mDia1, and mDia2 with microtubules. Mol Biol Cell. 2011;22(23):4575–4587.

27. Pimm ML, Henty-Ridilla JL. New twists in actin–microtubule interactions. Mol Biol Cell. 2021;32(3):211–217.

28. Innocenti M. New insights into the formation and the function of lamellipodia and ruffles in mesenchymal cell migration. Cell Adh Migr. 2018;12(5):401–416.

29. Sun H, Schlondorff J, Higgs HN, Pollak MR. Inverted formin 2 regulates actin dynamics by antagonizing Rho/diaphanous-related formin signaling. JASN. 2013;24(6):917–929.

30. Subramanian B, Chun J, Perez-Gill C, et al. FSGS-causing INF2 mutation impairs cleaved INF2 N-fragment functions in podocytes. JASN. 2020;31(2):374–391.

31. Yang C, Czech L, Gerboth S, Kojima S-i, Scita G, Svitkina T. Novel roles of formin mDia2 in lamellipodia and filopodia formation in motile cells. PLoS Biol. 2007;5(11):e317.

32. Krendel M, Pruyne D. New paradigm for cytoskeletal organization in podocytes: Proteolytic fragments of INF2 formin function independently of INF2 actin regulatory activity. JASN. 2020;31(2):235–236.

33. Salzer JL. Schwann cell myelination. CSH PERSPECT BIOL. 2015;7(8):a020529.

34. Mu A, Fung TS, Kettenbach AN, Chakrabarti R, Higgs HN. A complex containing lysine-acetylated actin inhibits the formin INF2. Nat Cell Biol. 2019;21(5):592–602.

35. Fernández-Barrera J, Alonso MA. Coordination of microtubule acetylation and the actin cytoskeleton by formins. Cell Mol Life Sci. 2018;75(17):3181–3191.

36. Ramabhadran V, Hatch AL, Higgs HNJJoBC. Actin monomers activate inverted formin 2 by competing with its autoinhibitory interaction. J Biol Chem. 2013;288(37):26847–26855.

37. Schneider R, Deutsch K, Hoeprich GJ, et al. DAAM2 variants cause nephrotic syndrome via actin dysregulation. Am J Hum Genet. 2020;107(6):1113–1128.

38. Jing J, Liu G, Huang Y, Zhou Y. A molecular toolbox for interrogation of membrane contact sites. J Physiol 2020;598(9):1725–1739.

39. Blackstone C. Cellular pathways of hereditary spastic paraplegia. Annu Rev Neurosci. 2012;35:25–47.

40. Friedman J, Webster B, Mastronarde D, et al. ER sliding dynamics and ER– mitochondrial contacts occur on acetylated microtubules. J Cell Biol. 2010;190(3):363–375.

41. Bartolini F, Andres-Delgado L, Qu X, et al. An mDia1-INF2 formin activation cascade facilitated by IQGAP1 regulates stable microtubules in migrating cells. J Mol Cell Biol. 2016;27(11):1797–1808.

42. Ishizaki T, Morishima Y, Okamoto M, Furuyashiki T, Kato T, Narumiya S. Coordination of microtubules and the actin cytoskeleton by the Rho effector mDia1. Nat Cell Biol. 2001;3(1):8–14.

43. Fung TS, Chakrabarti R, Higgs HN. The multiple links between actin and mitochondria. Nat Rev Mol Cell Biol. 2023;24(9):651–667.

44. Rogers SL, Gelfand VI. Membrane trafficking, organelle transport, and the cytoskeleton. COCEBI. 2000;12(1):57–62.

45. Kraus F, Roy K, Pucadyil TJ, Ryan MT. Function and regulation of the divisome for mitochondrial fission. Nature. 2021;590(7844):57-66.

46. Palau F, Estela A, Pla-Martín D, Sánchez-Piris M. The role of mitochondrial network dynamics in the pathogenesis of Charcot-Marie-Tooth disease. Adv Exp Med Biol. 2009:129–137.

47. Chan DC. Fusion and fission: interlinked processes critical for mitochondrial health. Annu Rev Genet. 2012;46:265–287.

48. Atkinson S, Doberstein S, Pollard T. Moving off the beaten track. Curr Biol. 1992;2(6):326–328.

49. Cabukusta B, Neefjes J. Mechanisms of lysosomal positioning and movement. Traffic. 2018;19(10):761–769.

50. Sun H, Perez-Gill C, Schlöndorff JS, Subramanian B, Pollak MR. Dysregulated dynein-mediated trafficking of nephrin causes INF2-related podocytopathy. JASN. 2021;32(2):307–322.

51. Maiti S, Michelot A, Gould C, Blanchoin L, Sokolova O, Goode BL. Structure and activity of full-length formin mDia1. Cytoskeleton. 2012;69(6):393–405.

52. Otomo T, Otomo C, Tomchick DR, Machius M, Rosen MK. Structural basis of Rho GTPase-mediated activation of the formin mDia1. Mol Cell. 2005;18(3):273–281.

53. Verderame M, Alcorta D, Egnor M, Smith K, Pollack R. Cytoskeletal F-actin patterns quantitated with fluorescein isothiocyanate-phalloidin in normal and transformed cells. PNAS. 1980;77(11):6624–6628.

54. Yamashiro S, Taniguchi D, Tanaka S, Kiuchi T, Vavylonis D, Watanabe N. Convection-induced biased distribution of actin probes in live cells. Biophys J. 2019;116(1):142–150.

55. Urbančič V, Butler R, Richier B, et al. Filopodyan: An open-source pipeline for the analysis of filopodia. J Cell Biol. 2017;216(10):3405–3422.

